# Temporally stable beta sensorimotor oscillations and cortico–muscular coupling underlie force steadiness

**DOI:** 10.1101/2021.11.30.470537

**Authors:** Scott J. Mongold, Harri Piitulainen, Thomas Legrand, Marc Vander Ghinst, Gilles Naeije, Veikko Jousmäki, Mathieu Bourguignon

## Abstract

As humans, we seamlessly hold objects in our hands, and may even lose consciousness of these objects. This phenomenon raises the unsettled question of the involvement of the cerebral cortex, the core area for voluntary motor control, in dynamically maintaining steady muscle force. To address this issue, we measured magnetoencephalographic brain activity from healthy adults who maintained a steady pinch grip. Using a novel analysis approach, we uncovered fine-grained temporal modulations in the ∼20-Hz sensorimotor brain rhythm and its coupling with muscle activity, with respect to several aspects of muscle force (rate of increase/decrease or plateauing high/low). These modulations preceded changes in force features by ∼40 ms and possessed behavioral relevance, as less salient or absent modulation predicted a more stable force output. These findings have consequences for the existing theories regarding the functional role of cortico-muscular coupling, and suggest that steady muscle contractions are characterized by a stable rather than fluttering involvement of the sensorimotor cortex.

## Introduction

As humans, we rely on our hands to interact with the environment, using them to communicate, touch, and importantly, hold items, i.e., phone, coffee, and keys. Remarkably, we are able to lose awareness of the very object in our hand, even as we maintain grip, begging the question: is voluntary control necessary for sustained, low intensity steady contractions? The sensorimotor cortex is unequivocally implicated in voluntary muscle contraction, yet, its role in maintaining steady contractions is unsettled. Does it play a sustained role in this highly dynamic process, or a phasic role in correcting when the goal is no longer matched (the phone is slipping off the hand)? At least in animals, stereotyped motor actions such as walking do not require corticomuscular communication after initiation (Purves, 1999).

When attempting to sustain an isometric contraction, the applied force is never constant but rather fluctuates around an average value (as reviewed in Enoka and Farina, 2021). Typically, force variability is quantified over an entire isometric contraction, usually maintained over several seconds (Laidlaw et al., 2000; Jones et al., 2002; Tracy and Enoka, 2002; Ushiyama et al., 2017). Surprisingly, there has been little investigation into non-global, ‘dynamic’ measures of force variability, and the neural mechanisms underlying regulation of force fluctuations have not been elucidated.

One means to assess corticomuscular communication is corticomuscular coherence (CMC). The literature suggests that CMC captures the phase coupling that occurs between brain and muscle activities, mainly at ∼20 Hz (Conway et al., 1995; Baker et al., 1997; Salenius et al., 1997) and ∼10 Hz (Piitulainen et al., 2015a), i.e, the main components of the sensorimotor rhythm that reflect the state of activation of sensorimotor cortices (Pineda, 2005). CMC usually peaks during sustained isometric contractions, decreases during dynamic contractions, and possesses somatotopic representation in the primary motor cortex and primary somatosensory cortex contralateral to the contracted muscle (Salenius et al., 1997; Hari and Salenius, 1999; Kilner et al., 1999; Salenius and Hari, 2003). A host of studies suggests it builds on the descending motor command (Bourguignon et al., 2019), but is modulated by (re)afferent information (Fisher et al., 2002; Kilner et al., 2004; Riddle and Baker, 2005; Liu et al., 2019).

Previous studies have focused on the association between global measures of force stability and CMC magnitude, based on minute-long recordings. Studies assessing motor precision where CMC levels are compared between conditions suggest that increased CMC is associated with smaller errors between target and exerted forces (Kristeva et al., 2007; Mendez-Balbuena et al., 2012). Somewhat contrastingly, in studies assessing correlation across participants, CMC appears positively associated with the amplitude of force fluctuations (Ushiyama et al., 2011a, 2017). However, no matter the reasons for the discrepancy, these studies did not look at the temporal dynamics of CMC in relation to force fluctuations, which is key to clarify the cortical involvement in force regulation.

Existing CMC analysis methods do not allow for the study of force regulation. CMC is typically estimated based on second-long epochs (Ushiyama et al., 2010, 2017; Mendez-Balbuena et al., 2012). This has allowed for assessment of CMC modulation in response to well-controlled isolated events, such as force ramps (Kilner et al., 2000, 2003) or sensory stimulations (Hari et al., 2014; Piitulainen et al., 2015b), but not continuously throughout a contraction. Here, we introduce a novel analysis method to identify the temporal dynamics of CMC and brain rhythms in relation to continuous signals. We use this method to determine how CMC and the ∼20-Hz brain rhythm modulate in relation to force fluctuations during volitional contraction. The main aim of the study is to determine if such modulations exist, arguing for a sustained role of the cortex in regulating steady contraction force. If so, we aim to determine (i) the temporal dynamics of these modulations with respect to force fluctuations, (ii) their relevance for force steadiness, and (iii) how they relate to global cortical involvement in the task (global CMC and ∼20-Hz power depression or enhancement).

## Materials and Methods

This is a reanalysis of previously published data (Bourguignon et al., 2017).

### Participants

Seventeen healthy human volunteers (7 females, 10 males; mean ± SD, age 34 ± 7 years, range 20–47 years) with no history of neuropsychiatric diseases or movement disorders participated in our study. All participants were right-handed (mean ± SD, score 90 ± 12, range 65–100 on the scale from –100 to 100; Edinburgh handedness inventory; Oldfield, 1971).

The study had prior approval by the ethics committee of the Helsinki and Uusimaa hospital district. The participants gave informed consent before participation, and they were compensated monetarily for travel expenses and lost working hours.

### Experimental protocol

Figure 1 illustrates the experimental paradigm. During MEG recordings, the participants were sitting with their left hand on the thigh and their right hand on a table in front of them. Participants’ vision was optically corrected with nonmagnetic goggles when needed. Participants were asked to maintain a steady isometric pinch grip of 2–4 N against a custom-made handgrip (connected to a rigid load cell; rigidity 15.4 N/mm; Model 1004, Vishay Precision Group, Malvern, PA, USA) with the right thumb and the index finger (see Fig. 1A), and to fixate at a black cross displayed on the center of a screen placed 1 m in front of them. When the force stepped out of the prescribed limits, a triangle (pointing up or down) appeared on top of the black cross, indicating in which direction to adjust the force, and it disappeared as soon as the force was correctly returned within the limits (see Fig. 1B and C). After a ∼1-min practice session, two 5-min blocks were recorded, with a minimum of 2-min rest between the blocks. Each block started with ∼10 s without contraction, after which participants were prompted to begin the contraction task. A 5-min task-free block was also recorded.

**Figure 1.**
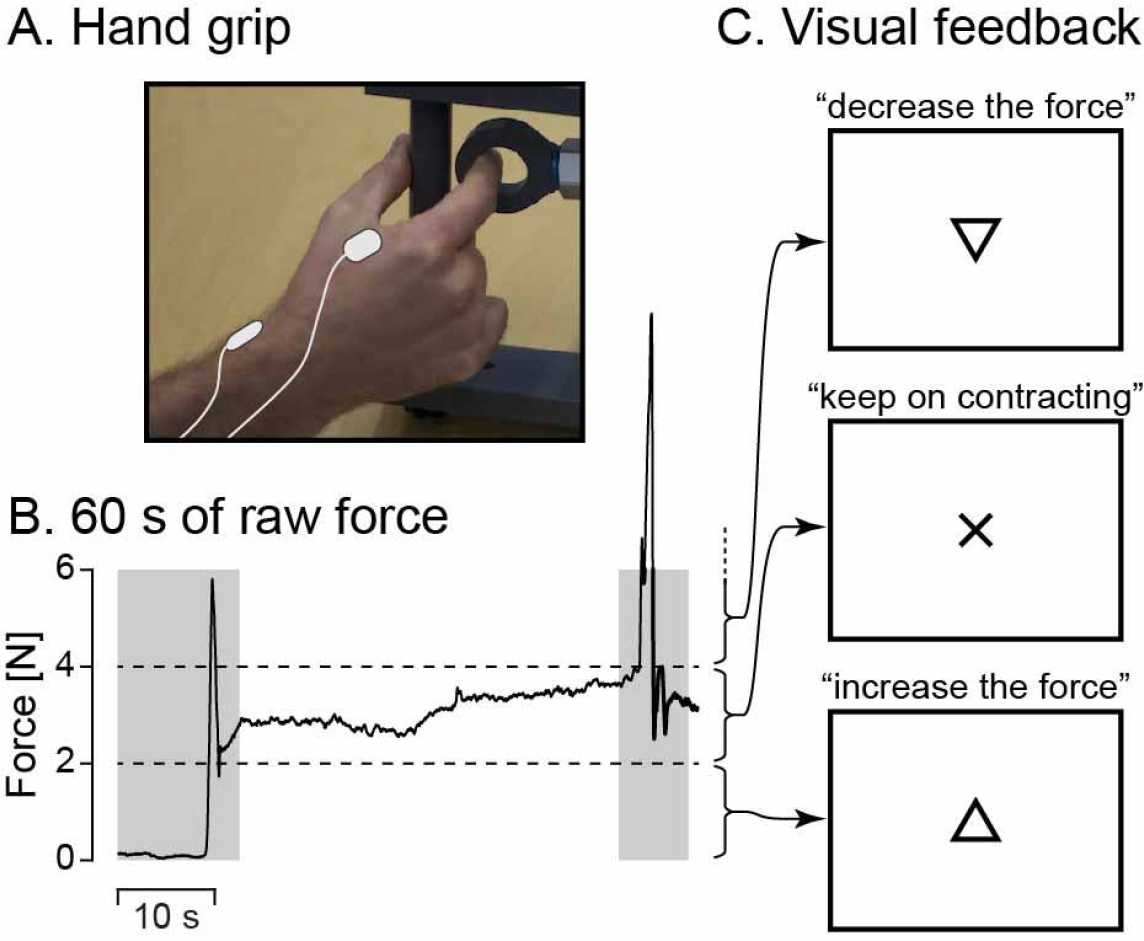
Experimental setup. A, Illustration of the isometric contraction task. A steady contraction is maintained on a custom-made handgrip with the right thumb and the index finger. Surface EMG is measured from the first dorsal interosseous (top right electrode) and the flexor carpi ulnaris (not visible here) of the right hand, with reference electrode over the distal radial bone (bottom left electrode). B, Sixty seconds of raw force signal from a representative participant. The participant was prompted to start contracting 10 s after the beginning of the recording. Two horizontal dashed lines indicate the force limits (2– 4 N). Gray shaded areas represent periods wherein contraction force was out of the prescribed bounds for at least 1 time-bin 2 s around. Corresponding data were not analyzed. C, Visual feedback presented to the participants to help them regulate their contraction force. Cross on the screen informed them that the force was within the prescribed limits. Arrow pointing up (respectively down) prompted them to increase (respectively decrease) the force. Reproduced from Bourguignon et al. (2017).

### Measurements

*MEG.* The MEG measurements were carried out in a magnetically shielded room (Imedco AG, Hägendorf, Switzerland) at the MEG Core of Aalto NeuroImaging, Aalto University, with a 306-channel whole-scalp neuromagnetometer (Elekta Neuromag™, Elekta Oy, Helsinki, Finland). The recording passband was 0.1–330 Hz and the signals were sampled at 1 kHz. Participants’ head position inside the MEG helmet was continuously monitored by feeding current into four head-tracking coils located on the scalp; the locations of the coils and at least 200 head-surface points (scalp and nose) with respect to anatomical fiducials were determined with an electromagnetic tracker (Fastrak, Polhemus, Colchester, VT, USA).

*EMG and force.* Surface EMG was measured from the *first dorsal interosseous*. Active EMG electrodes were placed on the muscle bulk and signals were measured with respect to a passive electrode placed over the distal radial bone. Recording passband was 10–330 Hz for EMG signals and DC–330 Hz for the force signal. EMG and force signals were then sampled at 1 kHz and recorded time-locked to MEG signals.

*MRI.* 3D-T1 magnetic resonance images (MRIs) were acquired with General Electric Signa^®^ 3.0 T whole-body MRI scanner (Signa VH/i, General Electric, Milwaukee, WI, USA) or with 3T MAGNETOM Skyra whole-body MRI scanner (Siemens Healthcare, Erlangen, Germany) at the AMI Centre, Aalto NeuroImaging, Aalto University School of Science.

### MEG preprocessing

Continuous MEG data were preprocessed off-line with MaxFilter 2.2.10 (Elekta Oy, Helsinki, Finland), including head movement compensation. The tSSS preprocessing was applied with a correlation limit of 0.9 and segment length equal to the recording length (Taulu and Kajola, 2005; Taulu and Simola, 2006). Independent component analysis was then applied to MEG signals filtered through 1–25 Hz, and 1–3 components corresponding to eye-blink and heartbeat artifacts were visually identified based on their topography and time-series. The corresponding components were subsequently subtracted from raw MEG signals.

### Conventional CMC estimation

We used coherence analysis to estimate CMC for all MEG sensors, and to identify the optimal MEG sensor for further analyses. Time points for which visual feedback was presented (force not properly kept between 2 and 4 N) were marked as bad to exclude periods during which contraction was intentionally corrected. Time points for which MEG signals exceeded 5 pT (magnetometers) or 1 pT/cm (gradiometers) were also marked as bad to avoid contamination of the data by any artifact not removed by the pre-processing. Continuous data from the recording blocks were split into 1000-ms epochs with 800-ms epoch overlap (Bortel and Sovka, 2007), leading to a frequency resolution of 1 Hz. Epochs less than 2-s away from time points marked as bad were discarded from further analyses (mean ± SD artifact-free epochs 2875 ± 130, range 2573–3031). Coherence spectra were computed between all MEG sensors and the non-rectified EMG signal following the formulation of Halliday et al. (1995), and by using the multitaper approach (5 orthogonal Slepian tapers, yielding a spectral smoothing of ± 2.5 Hz) to estimate power- and cross-spectra (Thomson, 1982). Data from gradiometer pairs were combined in the direction of maximum coherence as done in Bourguignon et al. (2015). Then, for each participant, the gradiometer pair with the highest coherence value in the 10–30-Hz band was selected among a predefined subset of 9 gradiometer pairs covering the primary sensorimotor (SM1) cortex. Further analyses were performed with the selected gradiometer signal in the orientation yielding the maximum coherence (MEG_SM1_).

### Force features

We extracted multiple features from the contraction force signal. First, the force signal was low-pass filtered at 30 Hz, as previously done in studies investigating force fluctuations during isometric contractions (Danion and Galléa, 2004; Witte et al., 2007; Missenard et al., 2009). Then, time points less than 2-s away from periods where the force changed over 50 ms by more than 5 SD (range for this threshold across participants, 0.13-0.35 N; mean ± SD, 0.25 ± 0.06) were marked as bad (and hence excluded from the analysis), completing those for which MEG amplitude was too high or the force was not in the prescribed range (2-4 N). From the low-pass filtered force signal *F*(*t*), we estimated the rate of force change as 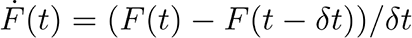 where is the time interval between adjacent samples (1 ms here). The two first force features considered were derived from this rate of force change (*Ḟ*):

1. The ***rate of force increase*** signal was the rate of force change half-wave rectified. That is *Ḟ* when *Ḟ* is positive and 0 otherwise.
2. The ***rate of force decrease*** signal was the opposite of the rate of force change half-wave rectified. That is - *Ḟ* when *Ḟ* is negative and 0 otherwise. The two additional force features considered were derived from the force plateauing signal, computed as the inverse of the rate of force change full-wave rectified 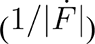. To render the inversion computationally stable, values of the force steadiness above the 95^th^ percentile were set to that percentile value. Of note, the choice of the clipping cutoff (percentile 90, 95 or 99) had virtually no impact on the results. These features were:
3. The ***force plateauing high*** signal was the force plateauing signal at time points where the force trace was concave (second derivative of *F,* 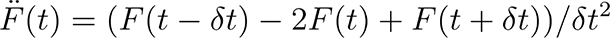, positive) and 0 elsewhere.
4. The ***force plateauing low*** signal was the force plateauing signal at time points where the force trace was convex (second derivative of *F* negative) and 0 elsewhere.

### CMC and power modulation in relation to force features

We next estimated how CMC and the power of MEG and EMG signals modulate in relation to the four force features. Key to this analysis is the use of Hilbert transformation to estimate coherence. In what comes next, we first present how the Hilbert transformation can be used to estimate coherence and power. We then generalize this approach to introduce a weighting by positive time-series, which we used to estimate CMC and power modulation in relation to the four selected force features presented in the previous subsection. Finally, we present the inclusion of time delays in force features to get at the temporal dynamics of CMC and power modulations.

To estimate CMC with the Hilbert transformation, MEG_SM1_ and EMG signals were filtered through a 5-Hz-wide frequency band centered on individual peak CMC frequency. The band-pass filter used in that effect was designed in the frequency domain with zero-phase and 1-Hz-wide squared-sine transitions from 0 to 1 and 1 to 0 (e.g. a filter centered on 20 Hz rose from 0 at 17 Hz to 1 at 18 Hz and ebbed from 1 at 22 Hz to 0 at 23 Hz). To remove the impact of artifacts, further analyses were based on time-points at least 2-s away from MEG artifacts (time points of MEG signals exceeding 5 pT for magnetometers or 1 pT/cm for gradiometers) and appearance of visual feedback. The analytical signals for MEG_SM1_ and EMG were then created by means of the Hilbert transform. From these analytical signals denoted *s*_1_(*t*) for MEG_SM1_ and *s*_2_(*t*) for EMG, the following formula

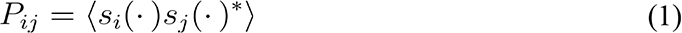

yielded MEG_SM1_ power (*i* = *j* = 1), EMG power (*i* = *j* = 2) and their cross-power (*i* = 1, *j* = 2). Here, the mean denoted ‹.› was taken across all artifact-free samples, and * denotes complex conjugation. From these quantities, the—unweighted—coherence is estimated as

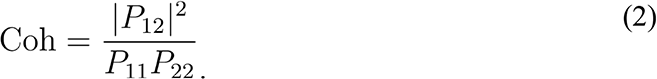

We used these formulas as a basis to introduce weighting by any positive time-series, which in our case will be each of the four force features. Denoting *w*(*t*) such a positive time-series time-locked to MEG and EMG signals, weighted auto- and cross-power can be estimated as

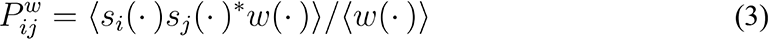

### and weighted coherence as

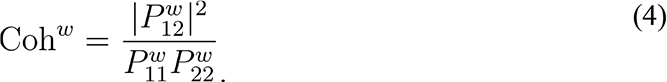

Notice that when *w* is constant, formulas (1) and (3) are equivalent. The interpretation of these weighted measures for non-constant *w* is straightforward. For example, an increase in Coh*^w^* compared with Coh indicates that CMC tends to be higher when the time-series considered (*w*) is high.

The last key element to our derivation of weighted coherence is the introduction of a time delay τ in *w*(*t*), so that 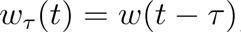 . From there, the temporal evolution of MEG_SM1_ and EMG power and their coherence with respect to *w* can be assessed as

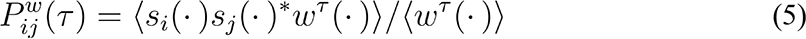

And

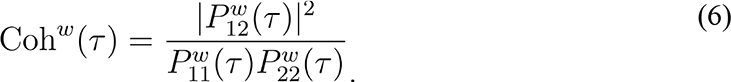

With these formulas, a peak increase in Coh*^w^* for a given force feature *w* at time delay, *e.g.*, = 40 ms, indicates that the force feature considered leads to an increase in CMC 40 ms later. Conversely, a peak in Coh*^w^* at a negative time delay indicates that high CMC leads to an increase in the force feature considered with time delay .

The analysis approach described above was used to estimate how ∼20-Hz CMC, EMG power, and MEG_SM1_ power modulate in relation to the four selected force features. In that analysis modulations were computed for time delays ( ) ranging from –2 s to 2 s by steps of 10 ms. Power modulations were normalized by the mean across the baseline defined by 1 s < | | < 2s. We also used this approach to estimate how full-band EMG power modulates in relation to the four force features. Full-band EMG power was obtained from the EMG signal high-pass filtered at 30 Hz, rectified, low-pass filtered at 50 Hz, and then squared. In that analysis, the temporal resolution ( ) was set to 1 ms to resolve the fast transient dynamics of EMG-force coupling.

### Link between identified modulations and behavioral relevance

We used cross-participant Pearson correlation to determine the degree of (in)dependence between and among the different modulations we observed, parameters quantifying force steadiness, global MEG_SM1_ power suppression, and static CMC values. These different measures are described below.

From CMC and power modulations, we extracted the maximum (for increases) or minimum (for decreases) value for each individual within ±50 ms around group-level peak values. These values were then converted to percentage of change relative to baseline (1 s < | | < 2), giving rise to CMC and power modulation values. For regularization purposes, peak and baseline CMC values were incremented by 0.01 before the division.

Force steadiness was estimated as the standard deviation (SD) of the 10-min of force signal filtered in three relevant frequency ranges: 0.5–3 Hz, 3–15 Hz and 15–30 Hz. Force fluctuations at 0.5–3 Hz were previously argued to be the most relevant for the maintenance of a steady contraction and to be monitored by the brain through the proprioceptive system (Bourguignon et al., 2017). Force fluctuations in the intermediate range (3–15 Hz) should relate more to intermittent motor control (Gross et al., 2002) and capture the physiological tremor at ∼10 Hz (McAuley et al., 1997; Gilbertson, 2005). Force fluctuations at 15–30 Hz were shown to be tightly linked to the presence of ∼20-Hz CMC (Bourguignon et al., 2017).

Global MEG_SM1_ power suppression was estimated as the MEG_SM1_ power at the frequency of peak CMC in the isometric contraction task divided by this same quantity estimated from the task-free recording (preprocessed in the same way as isometric contraction MEG data).

In cross-participant Pearson correlation, data points were considered as outliers if they departed by over 2 SDs from the mean (z-score above 2). A maximum of one outlier was left out (the one with the highest z-score) in each correlation analysis.

The levels of correlation between force SD and (*i*) dynamic CMC modulations and (*ii*) static CMC values were compared with the Steiger test (Steiger, 1980).

### Statistical analyses

#### Significance of unweighted coherence

A threshold for statistical significance of the coherence (*p* < 0.05 corrected for multiple comparisons) was obtained as the 95^th^ percentile of the distribution of the maximum coherence—across 10–30 Hz, and across the selection of 9 gradiometers—evaluated between MEG and Fourier-transform surrogate reference signals (1000 repetitions; Faes et al., 2004). The Fourier-transform surrogate of a signal is obtained by computing its Fourier transform, replacing the phase of the Fourier coefficients by random numbers in the range [−π; π], and then computing the inverse Fourier-transform (Theiler et al., 1992; Faes et al., 2004).

#### Significance of CMC and power modulation by force features

For each participant and each force feature, we used surrogate-data-based statistics to estimate the significance of the modulation in the 3 following signals: ∼20-Hz CMC, ∼20-Hz MEG_SM1_ power, and ∼20-Hz EMG power. Modulation amplitude was defined as the maximum difference in absolute value between modulation in the range -1 s to 1 s (| | < 1 s) and mean baseline value (1 s < | | < 2 s). A threshold for statistical significance of modulation amplitude (*p* < 0.05) was obtained as the 95^th^ percentile of its distribution for surrogate force features (1000 repetitions). Surrogate force features were derived from the genuine force feature by randomly shuffling the periods of non-zero force feature, and randomly shuffling the periods of zeros force feature.

Non-parametric permutation statistics were further used to estimate the significance at the group level of modulations by force features (Nichols and Holmes, 2002). This analysis aimed at determining if the modulation of each of the 3 modulated signals by each of the four force features was consistent across participants. First, we estimated the difference between the modulation signal averaged across participants for the genuine data, and for the first set of surrogate force features. We then built a permutation distribution (1000 permutations) for the maximum absolute value of such difference obtained after randomly swapping genuine and surrogate data for each participant. An exact *p*-value for each sample of the non-permuted data was obtained as the proportion of the values in the permutation distribution exceeding the value at this sample (Nichols and Holmes, 2002). Time points corresponding to significant modulations (*p* < 0.05) were identified. The permutation procedure was repeated 20 times, for each of the 20 first sets of surrogate modulation signals, and we report the mean values to smooth out estimation inaccuracies pertaining to the random character of the surrogate data.

#### Confidence interval for increases and decreases in CMC and power modulations

We used bootstrap statistics (Efron, 1979; Efron and Tibshirani, 1994) to determine a 95% confidence interval for the timing and amplitude of peaks and troughs of modulations ∼20-Hz CMC, ∼20-Hz MEG_SM1_ power, ∼20-Hz EMG power, and full-band EMG power in relation to force features. The bootstrap applied was bias-corrected and accelerated with 1000 resamplings (Efron, 1979; Efron and Tibshirani, 1994). It was applied to the time courses upsampled by a factor 10 using spline interpolation (i.e., 0.1 ms resolution for EMG full band, and 1 ms resolution for the 3 other modulation signals).

#### Global significance of associations with multiple modulation values

Modulations in ∼20-Hz CMC and MEG_SM1_ power were revealed to be highly correlated. Hence, we jointly estimated the significance of their correlation with other parameters (e.g., force SD or global MEG_SM1_ power attenuation) using non-parametric permutation statistics. Individual correlation values were averaged after having rendered their signs consistent (i.e., modulations were multiplied by -1 when they showed a negative correlation with an arbitrary reference modulation, the choice of which had no impact on the results). A permutation distribution was built for the averaged correlation value by computing 10,000 times its value after having randomly shuffled the values for the other parameter. The significance level was estimated as the proportion of values in the permutation distribution that are higher than the value for non-permuted data.

### Data availability

The code that supports the findings of this study is available on the Open Science Framework at (link that will be provided upon successful reviewing). The underlying numerical data for each figure can also be found in the supporting data files (which will be provided upon successful reviewing). Local ethical restrictions prevent us from sharing the raw data files.

## Results

### Static CMC

Figure 2 (left side) presents the spectra of CMC in all participants. CMC (mean ± SD, 0.058 ± 0.048) was statistically significant in 16/17 participants (*p* < 0.001) and marginally significant in the remaining participant (*p* = 0.060). It should be noted that the peak frequency and to a larger extent the magnitude of CMC were variable across participants, in line with previous studies (Ushiyama et al., 2011b).

**Figure 2.**
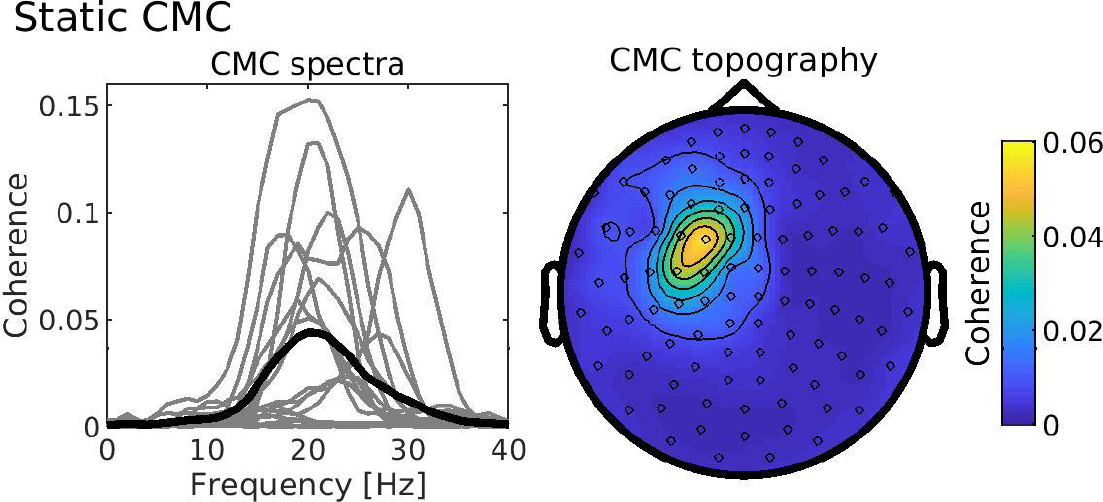
Static CMC. Left side — CMC spectra for each participant (gray traces) and group-average (black trace). For each participant, the trace corresponds to the CMC measured at the sensor (among the 9 sensors overlying the left SM1 cortex) for which CMC was maximum in the 10–30-Hz range. Right side — Spatial distribution of the CMC averaged across participants. CMC peaked above the left SM1 cortex.

Figure 2 (right side) shows the distribution of ∼20-Hz CMC in a representative participant. As expected, CMC was maximal in sensors above the left SM1 cortex.

### Validity of force features

Figure 3 presents a 500-ms excerpt of force signal and related force features. The force trace presents clear fluctuations of the order of 0.1 N. These fluctuations are well characterized by the four force features we selected. Clearly, the force features we extracted identify the short-lived events they were designed to capture.

**Figure 3.**
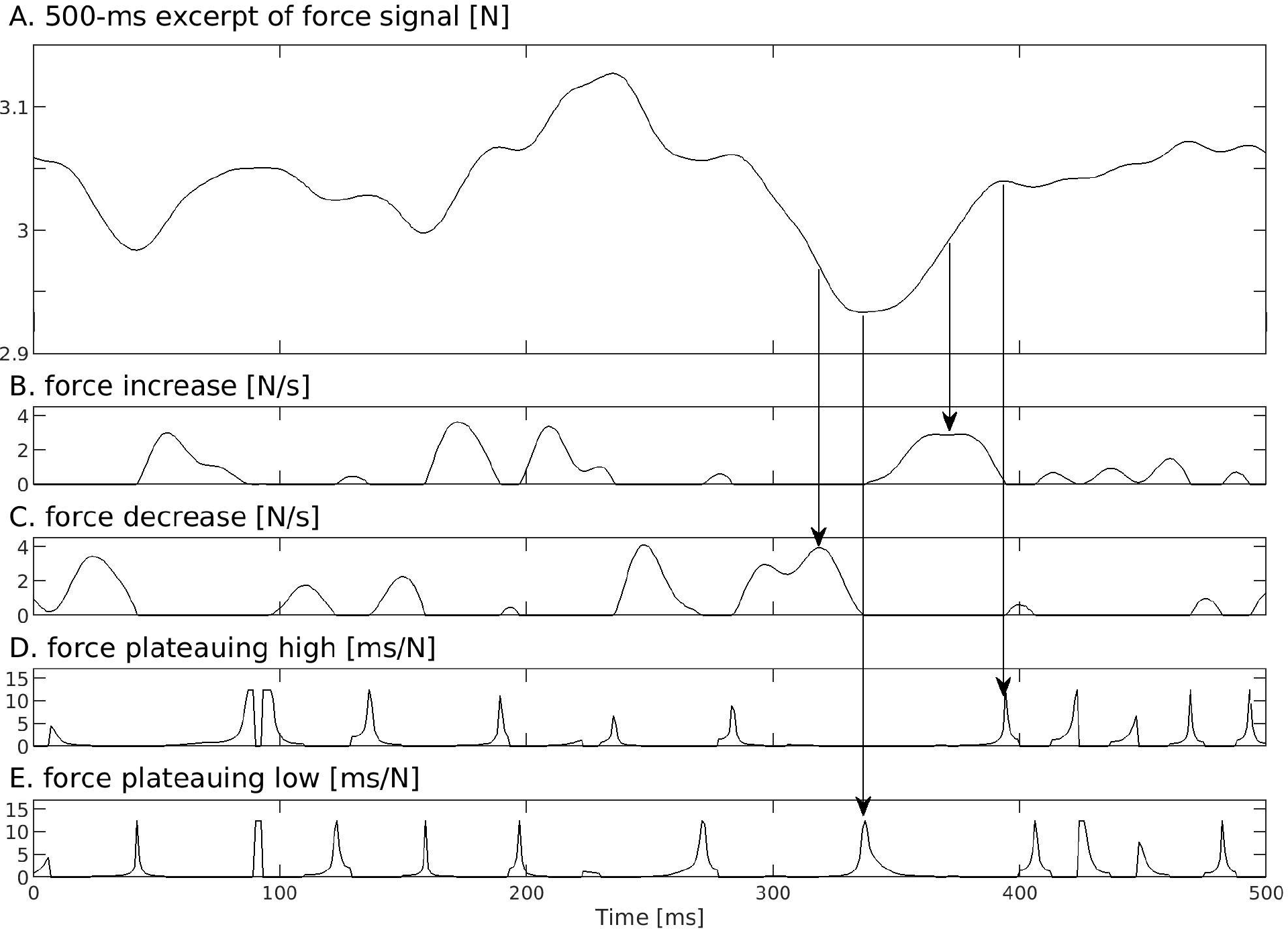
Excerpt (500 ms) of force signal (A) and related force features (B–E) for a representative participant. Vertical arrows indicate the correspondence between force features seen in the raw force signal and peaks in the force feature signals.

Figure 4 presents the auto- (Figure 4A) and cross-correlation (Figure 4B) of some of the force features. Force features were characterized by oscillations in their level of auto- and cross-correlations that essentially vanished within 200 ms of delays (| |). The period of the oscillations in auto-correlation was about 70 ms for force increase and decrease, and 20 ms for force plateauing high and low. This indicates that CMC and power modulations should not display side peaks attributable to auto-correlation in force features. This is because the intrinsic temporal granularity (or temporal resolution) of CMC and power (about 200 ms since their computation was based on signals restricted to a 5-Hz band) is lower (i.e., higher duration) than that of the auto-correlation (20–70 ms).

**Figure 4.**
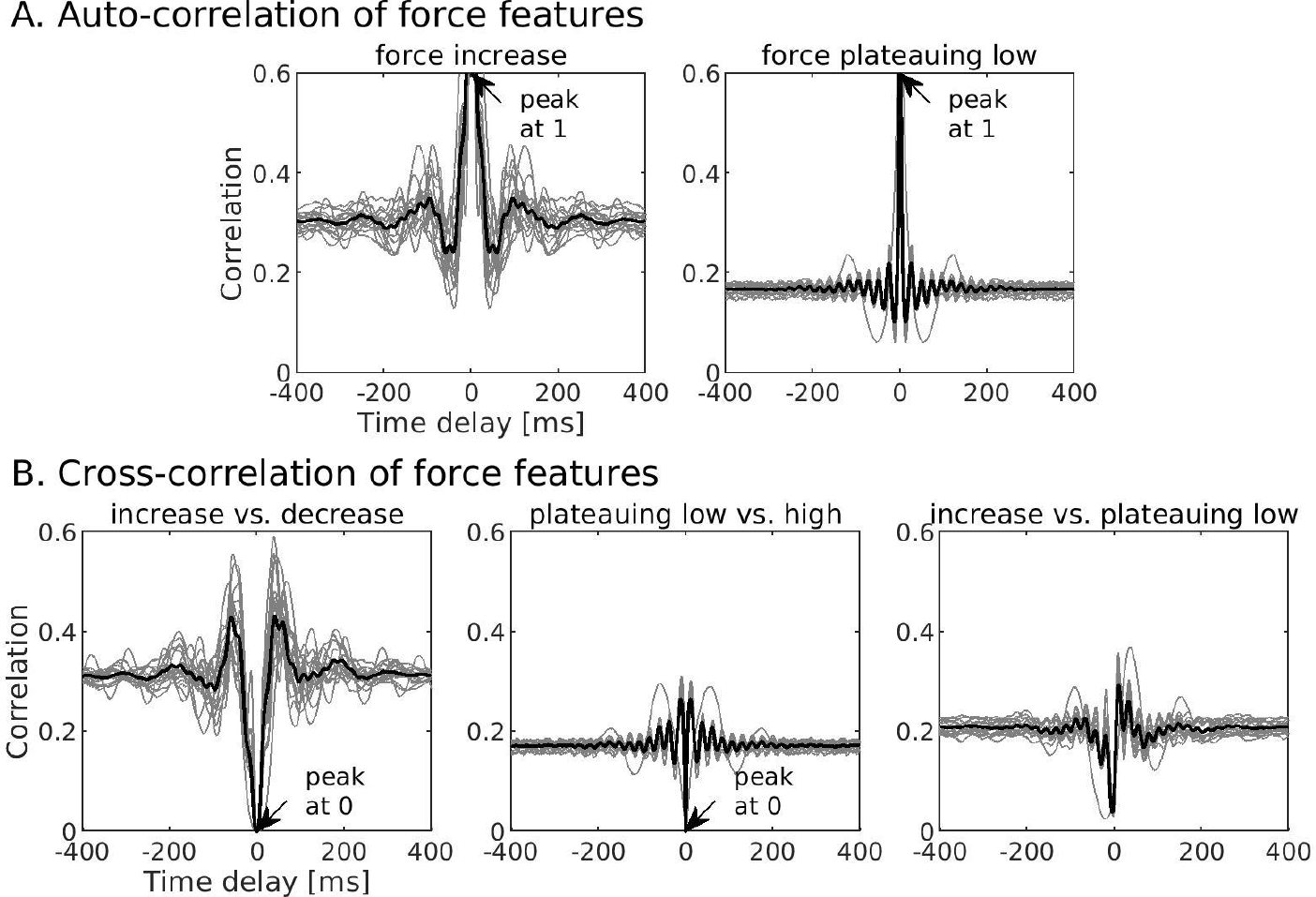
Auto-correlation (A) and cross-correlation (B) of force features. Traces are in gray for every participant and in back for the group-average. Auto-correlation for force decrease and plateauing high were highly similar to those for force increase and plateauing low (respectively). Cross-correlation between force increase and plateauing low were highly similar to those for the 3 other pairs involving increase/decrease on the one hand and plateauing high/low on the other hand.

Beyond delays of 200 ms, correlation reached a non-zero baseline level 0.2–0.3. The fact that baseline correlation was not 0 is attributable to the non-normality of the force feature signals (Bishara and Hittner, 2015).

Trivially, when the delay was 0 ( = 0), auto-correlations were 1 for all force features. For such null delay, the cross-correlation was 0 between force increase and decrease, and between force plateauing high and low. Such a value was expected because the first signal of each of these two pairs was non-zero only when the other was at zero and vice versa.

Beyond the initial peaks at = 0, the level of auto- and cross-correlation correlation was limited, with maximum values of about 0.1 (up to 0.3 for some participants) above baseline level. This indicates that force features were well suited to identify transient events that are not too interdependent.

In summary, the limited interdependence of force features and their fast-decaying auto-correlation ensure adequacy for modulatory analysis of CMC and power in relation to transient events during continuous isometric contraction.

### Physiological relevance of force features

To determine the physiological relevance of the identified force features and validate the novel analysis approach, we first focused on the modulation of wide-band EMG power in relation to the force features.

Figure 5 presents these modulations for each participant and averaged across participants. All participants displayed a significant modulation in wide-band EMG power in relation to all four force features (*p* < 0.003). In relation to the force increase signal, EMG power showed a peak at –13 ms followed by a trough at 8 ms (see Fig. 5 for confidence intervals and power modulation values). The opposite response pattern was observed for the force decrease signal. In relation to force plateauing high, EMG power showed a trough at –2 ms; the reverse pattern was seen in response to force plateauing low.

**Figure 5.**
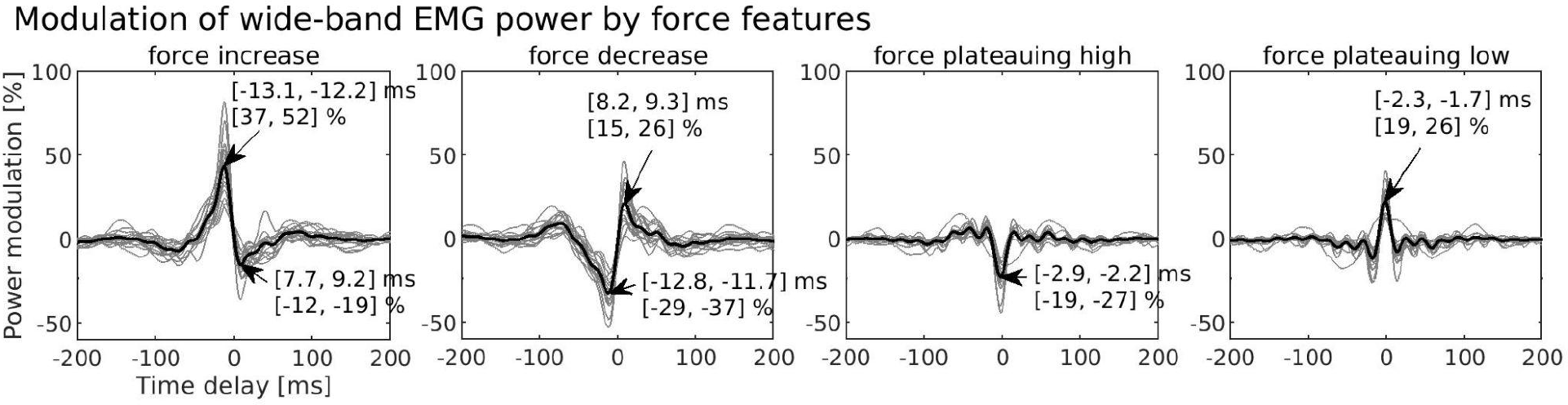
Modulation of wide-band (30–330 Hz) EMG power in relation to force features. From left to right, plots are for force increase, force decrease, force plateauing high, and force plateauing low. Traces are in gray for every participant and in back for the group-average. Power modulation is expressed in percentage of change relative to baseline (mean across 1 s < | | < 2 s), as function of the time delay ( ) with respect to the considered force feature. Arrows indicate the location of peaks and troughs as well as a bootstrap confidence interval for their timing and modulation amplitude.

### Modulation of ∼20-Hz CMC, MEG_SM1_ power and EMG power in relation to force features

Figure 6 presents the modulation of ∼20-Hz CMC, ∼20-Hz MEG_SM1_ power and ∼20-Hz EMG power in relation to the four investigated force features.

**Figure 6.**
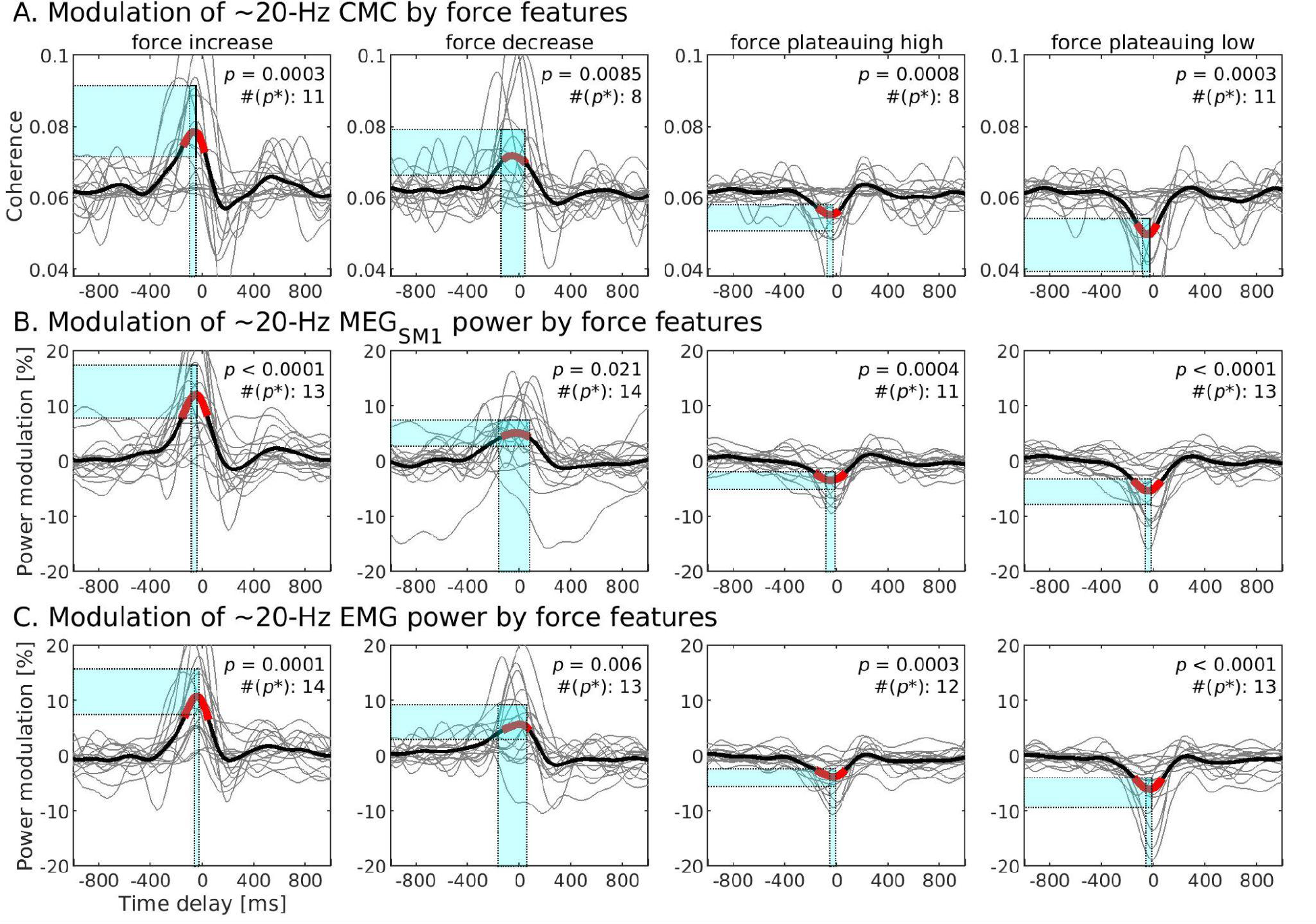
Modulation of ∼20-Hz CMC (A), ∼20-Hz MEG_SM1_ power (B) and ∼20-Hz EMG power (C) in relation to force features. Traces are in gray for every participant and in black for the group-average. The time intervals associated with significant modulation (p < 0.05 corrected for multiple comparisons; permutation statistics) are highlighted in red; exact p-values are provided in the top-right corners, along with the number of participants showing statistically significant modulation (p < 0.05; surrogate-data-based statistics). Bootstrap confidence intervals for the peak values are indicated with cyan shaded areas. Individual coherence traces (part A) were translated vertically so their baseline value (mean across 1 s < |t| < 2 s) aligns with that of the group-average. Power modulation is expressed in percentage of change relative to baseline.

Overall, these three ∼20-Hz modulations displayed significant modulations in relation to the four force features. These modulations were statistically significant (*p* < 0.05) in at least half of the participants (see Figure 6 for exact numbers). The three measures showed a peak in relation to *force increase* and *force decrease*, and a trough in relation to *force plateauing high* and *force plateauing low*. All peaks or troughs were consistent across participants (i.e., in the same direction and with similar latency), so that they were significant at the group level (*p* < 0.05 in all 12 instances; see Figure 6 for exact *p*-values). In all instances, the latency of the modulation tended to precede the force features by ∼40 ms (see Figure 6 for bootstrap confidence intervals).

Some force features induced stronger modulations than others. Peaks of ∼20-Hz CMC were higher in relation to *force increase* than *decrease* (27.6% vs. 17.5%, *t*_16_ = 2.37, *p* = 0.012); the same occurred also for ∼20-Hz MEG_SM1_ power (12.0% vs. 5.1%, *t*_16_ = 3.66, *p* = 0.0021) and ∼20-Hz EMG power (10.7% vs. 5.6%, *t*_16_ = 2.62, *p* = 0.018). Also, the suppression of ∼20-Hz CMC was more pronounced in relation to *force plateauing low* compared to *high* (–19.5% vs. –10.5%; *t*_16_ = 3.46, *p* = 0.0067); the same occurred also for ∼20-Hz MEG_SM1_ power (–5.3% vs. –3.5%, *t*_16_ = 3.69, *p* = 0.0020) and ∼20-Hz EMG power (–6.0% vs. –3.8%, *t*_16_ = 3.87, *p* = 0.0014). Importantly, *force plateauing low* gave rise to stronger CMC depression in all participants showing significant modulation in relation to either of the force plateauing signals. Moreover, these significant effects evidenced by group-level comparisons were well mirrored by individual data. Among the participants showing significant modulations in response to either *force increase* or *decrease*, the modulation in relation to *force increase* was higher in 10/13 participants for CMC, 13/15 for ∼20-Hz MEG_SM1_ power, and 12/15 for ∼20-Hz EMG power. Among the participants showing significant modulations in relation to either *force plateauing low* or *high*, the modulation was stronger in relation to *force plateauing low* in 13/13 participants for CMC, 13/15 for ∼20-Hz MEG_SM1_ power, and 13/13 for ∼20-Hz EMG power. In light of this, further analyses will be conducted only on modulations in relation to *force increase* and *force plateauing low*.

Given that ∼20 Hz CMC and power present modulations in the same direction (increases in relation to force increase/decrease; decreases in relation to force plateauing high/low), it is not possible to tell whether CMC modulations are attributable to changes in the signal-to-noise ratio in the MEG_SM1_ and EMG signals. In fact, the modulations we observed were related in multiple ways. Figure 7 shows the existence of strong associations between ∼20-Hz CMC and MEG_SM1_ power modulations (Fig. 7A), and between each of these two measures assessed in relation to *force increase* versus *force plateauing low* (Fig. 7B).

**Figure 7.**
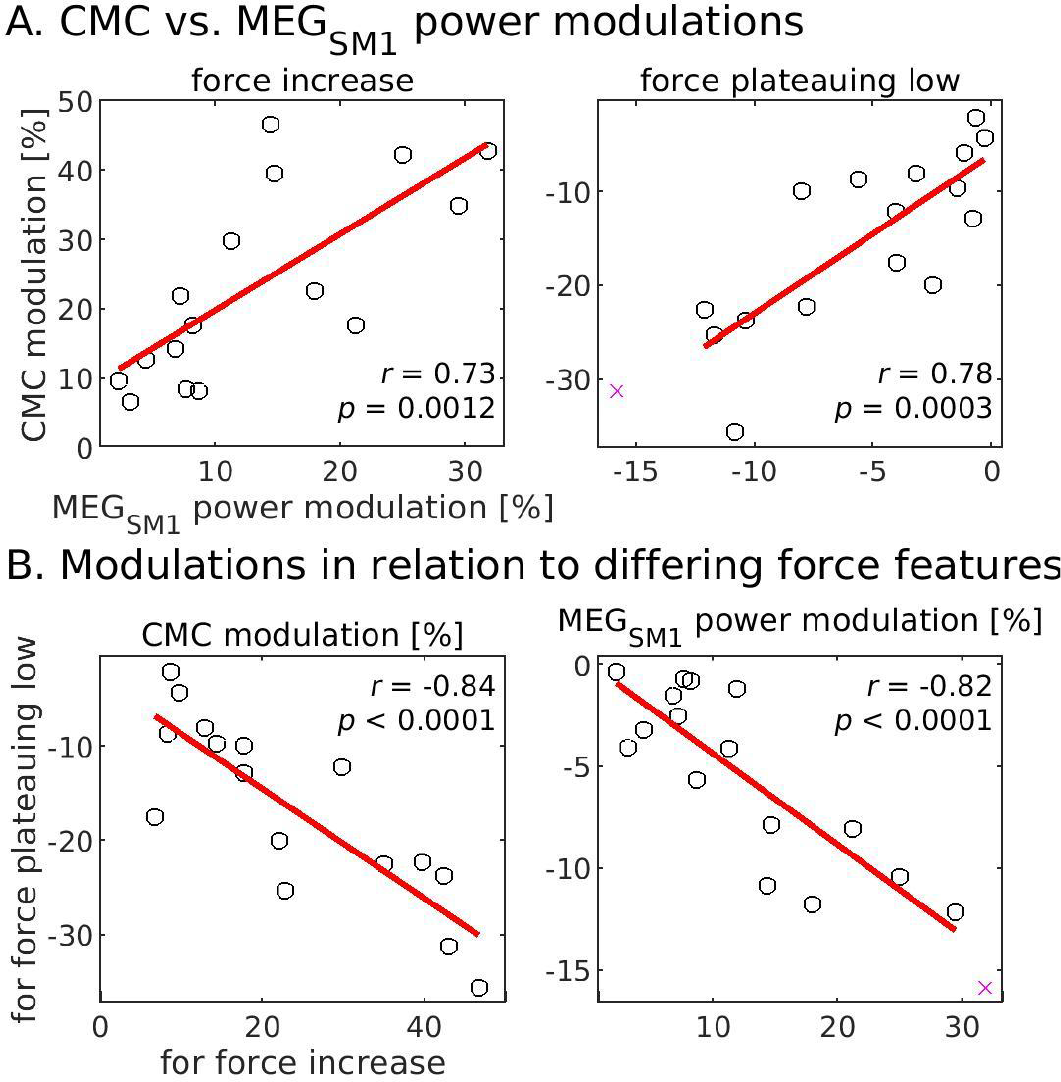
Interdependence between uncovered modulations. A — Relation between ∼20-Hz CMC and MEG_SM1_ power moduations in relation to force increase (left) and force plateauing low (right). B — Relation between modulations in relation to force plateauing low vs. force increase for ∼20-Hz CMC (left) and ∼20-Hz MEG_SM1_ power. In each graph, black circles indicate individual values included in the correlation analysis, red crosses indicate outliers (potentially different participants in the different graphs), a red line was obtained by linear regression, and correlation values with associated significance level are indicated in the top or bottom right corner.

### Relevance of identified ∼20-Hz modulations for force steadiness

We next determine the relevance for force steadiness of the most salient ∼20-Hz modulations in CMC and MEG_SM1_, in relation to *force increase* and *force plateauing low*. For that, we sought a linear association between individual values of CMC or power increase/decrease and force SD in different frequency ranges (0.5–3 Hz, 3–15 Hz and 15–30 Hz). The unique global permutation test revealed a significant association between modulations—in ∼20-Hz CMC and MEG_SM1_ power in relation to *force increase* and *force plateauing low*—and force SD at 0.5–3 Hz (*p* = 0.043), 3–15 Hz (*p* = 0.042) and 15–30 Hz (*p* =

0.0024).

Figure 8 presents the associations for *force plateauing low*. There was a significant negative correlation between force SD in all tested frequency ranges and the modulation in both

**Figure 8.**
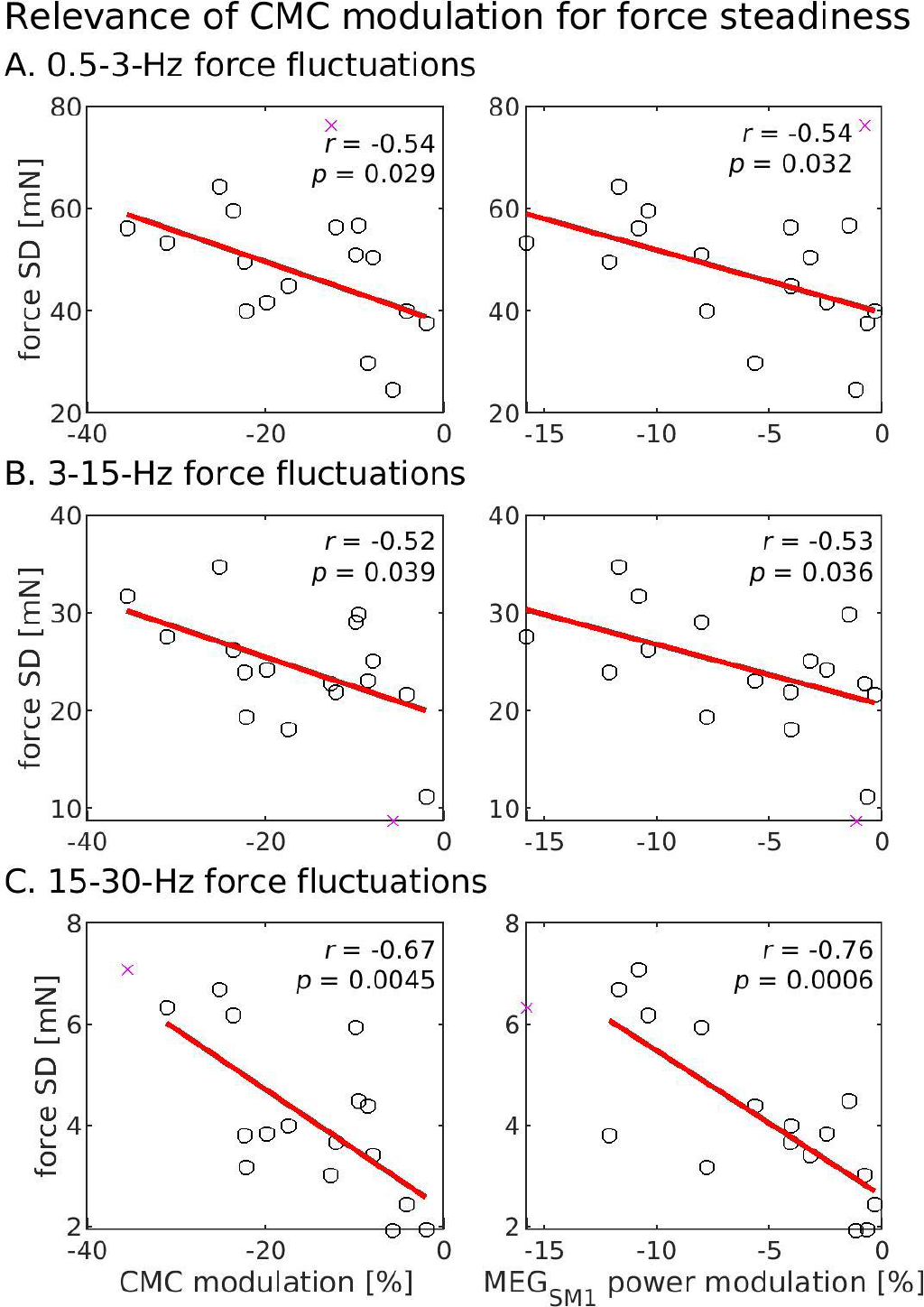
Relevance for force steadiness of the modulations identified in relation to force plateauing low. Each graph presents the SD of the raw force signal filtered through 0.5–3-Hz (A), 3–15-Hz (B) and 15–30-Hz (C) as function of the percentage of change in ∼20-Hz CMC (left) and MEG_SM1_ power (right) at the trough of the modulation relative to baseline. Refer to Figure 7 for a description of graph layout.

∼20-Hz CMC and MEG_SM1_ power. The association was especially salient with force SD at 15–30 Hz. Associations for the *force increase* signal were all positive but were significant only for the force SD at 15–30 Hz (*r* = 0.58–0.62, *p* = 0.011–0.018) and not in the two other frequency ranges (*r* = 0.31–0.40, *p* = 0.12–0.24).

### Relation between identified ∼20-Hz modulations and global ∼20-Hz power depression

We next asked if inter-individual variability in the amplitude of the observed dynamic modulation in ∼20-Hz CMC and MEG_SM1_ power relates to inter-individual variability in global depression in the ∼20-Hz MEG_SM1_ power during contraction compared with task-free power.

Specifically, we aimed to determine if weaker dynamic modulations reflect a state where the ∼20-Hz sensorimotor rhythm is (*i*) unaffected by the contraction task or (*ii*) continuously attenuated.

A unique global test revealed a non-significant trend of association between modulations—in ∼20-Hz CMC and MEG_SM1_ power in relation to *force increase* and *force plateauing low*—and the ratio between global ∼20-Hz power during isometric contraction and while not performing the task (*p* = 0.083; non parametric permutation test).

Figure 9 presents the associations for the modulations in relation to *force plateauing low*. Although the associations were not significant, they were more in line with the second option: the weaker the dynamic modulation (in ∼20-Hz CMC and MEG_SM1_ power), the deeper the global depression in ∼20-Hz MEG_SM1_ power.

**Figure 9.**
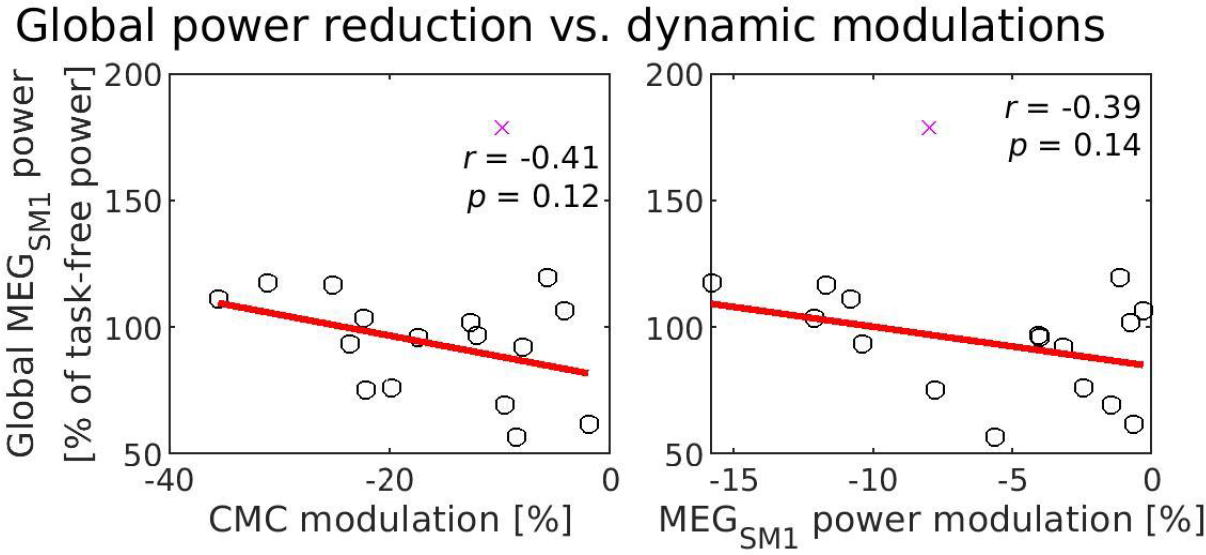
Relation between global ∼20-Hz MEG_SM1_ power during isometric contraction (expressed in percentage of power in the task-free condition) and modulation in CMC (left) and power (right) in relation to force plateauing low. All is as in Figure 7.

### Relation between static CMC and dynamic CMC modulations

We next asked whether dynamic modulations in CMC are linked to the global (static) level of CMC. In other words, do individuals with stronger levels of static CMC present more prominent dynamic modulations?

A unique global test revealed no significant association between modulations—in ∼20-Hz CMC in relation to *force increase* and *force plateauing low*—and the static values of ∼20-Hz CMC (*p* = 0.26). The scatter plots did not highlight the presence of a trend for either of the two force features (Fig. 10). Also worth remembering, the regularization procedure used to estimate CMC modulation was to the effect of reducing modulation values for participants with CMC values not well above 0.01. Hence, the correlation was naturally biased to negative values, but still, did not come significant.

**Figure 10.**
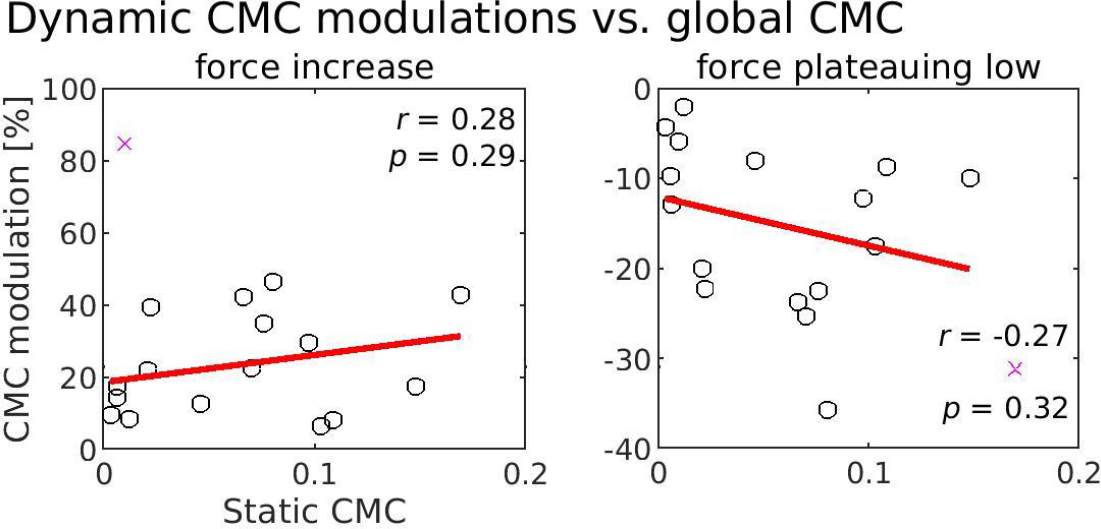
Relation between static CMC and dynamic CMC modulation in relation to force increase (left) and force plateauing low (right).

Since static CMC and dynamic CMC modulations are essentially decoupled, we asked if static CMC is relevant for the maintenance of the static contraction, and if it is more relevant than dynamic CMC modulations.

Figure 11 shows that the level of static CMC correlates significantly with the SD of 15–30 Hz force fluctuations, but not of 0.5–3 Hz and 3–15 Hz force fluctuations. However, the correlation with static CMC was not significantly different from that with CMC modulation for any of the three frequency bands (*p* > 0.2; Steiger test).

**Figure 11.**
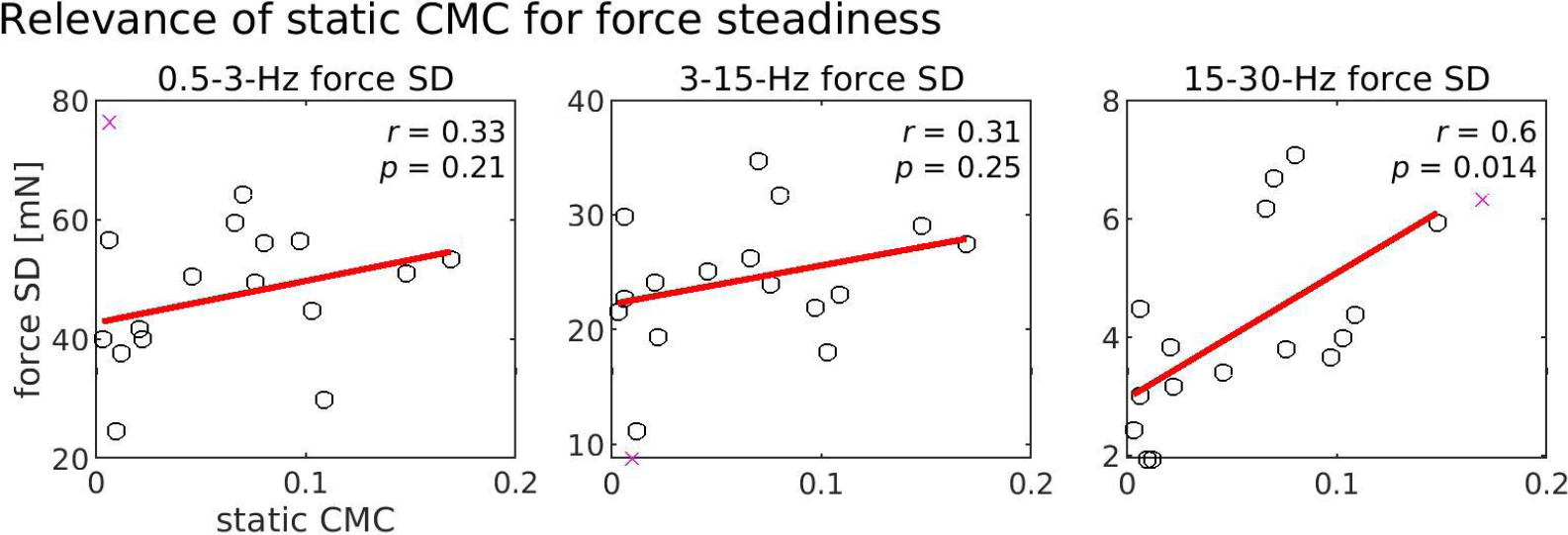
Relevance of static CMC for the steadiness of the contraction force at 0.5–3 Hz (A), 3–15 Hz (B) and 15–30 Hz (C).

## Discussion

In the present study, we examined how ∼20-Hz CMC, MEG_SM1_ power, and EMG power modulate in relation to different force features (rate of force change and plateaus) during submaximal isometric contractions. We found consistent temporal modulations that preceded changes in force features by ∼40 ms. The amplitude of these modulations, reflecting the extent of fluctuations in cortical involvement, was associated with force variability, which indicates behavioral relevance. These results are key to understanding the role of the SM1 cortex in dynamically maintaining steady muscle force.

### Steady contraction entails stable ∼20-Hz SM1 oscillations and CMC

All identified modulations in ∼20-Hz CMC were accompanied by concomitant power modulations at ∼20-Hz in MEG_SM1_ and EMG signals. The same observation was made in previous studies of CMC and power modulations in relation to discrete events (Kilner et al., 2000, 2003; Hari et al., 2014; Piitulainen et al., 2015b). In such context, it is unclear whether a change in CMC pertains to a genuine change in phase locking, or to a change in the amplitude of the coherent signal, which leads to increased coherence estimation via increased signal-to-noise ratio (Muthukumaraswamy and Singh, 2011; Bayraktaroglu et al., 2013). Additionally, we identified interdependence of modulations (Fig. 7), where the magnitude of CMC and MEG_SM1_ power modulations for different force fluctuation signals shared a highly similar time-course. This means that CMC modulations could be attributable to changes in the signal-to-noise ratio in the MEG_SM1_ and EMG signals. Therefore, we will refrain from interpreting our results as evidence for dynamic changes in cortico-muscular communication driving force fluctuations.

There is a general consensus that neural synchronization, which gives rise to a MEG power increase in population-level cortical activity, is a suboptimal form of neural coding, in that it results in a reduction in degrees of freedom and processing of information content (Aumann and Prut, 2015). On the other hand, neural desynchronization, and associated decrease in MEG power, reflects more complex computation (Averbeck and Lee, 2004). Considering this computational view of cortical processing and a large body of empirical data, a reduction in the level of ∼20-Hz power is considered to reflect increased activation or involvement of the sensorimotor cortex (Pineda, 2005; Démas et al., 2019). With this mechanism in mind, our data suggests that SM1 cortex intermittently adjusts the level of corticospinal drive to reverse the direction of force change (reduction in ∼20-Hz power leads to force plateauing) to maintain steady contraction. In between successive adjustments, the SM1 cortex disengages or is inhibited through peripheral afference or other cortical and subcortical processes (increased ∼20-Hz power), leading to a maintenance of the motor state and associated corticospinal drive, and depending on the drive, to an increase or decrease of the force. Importantly, we found that individuals with lower modulations in ∼20-Hz power and CMC are able to maintain more steady contractions. Thus, low modulations in ∼20-Hz power and CMC would indicate a stable (rather than fluttering) state of SM1 cortex with regard to the contraction.

Our data does not provide a definite answer as to whether a stable state reflects constant cortical engagement or disengagement. However, a non-significant trend indicated that a higher attenuation of global ∼20-Hz power (taken as a percentage of task-free power) is associated with lower modulations in ∼20-Hz CMC and power (Fig. 9). This trend suggests that individuals who do not show the ∼20-Hz modulations have a tendency to continuously desynchronize their ∼20-Hz SM1 activity. Hence, stable contractions would involve sustained engagement of the SM1 cortex in order to regulate the contraction.

### Compatibility with hypothesized functional role of CMC

Three main hypotheses have been formulated regarding the functional role played by CMC in sensorimotor control. The *motor state maintenance* hypothesis posits that ∼20-Hz oscillations underlying CMC promote the maintenance of a stable motor state (Gilbertson, 2005; Androulidakis et al., 2007; Baker, 2007; Engel and Fries, 2010). The *test-pulse* hypothesis posits that CMC reflects the mechanism by which the sensorimotor system sends pulses to muscles at ∼20 Hz and monitors the resulting afferent signal to probe the state of the periphery for continuous sensorimotor recalibration (Baker, 2007; Mackay, 1997; Witham et al., 2011). The *SM1 rhythm corollary* hypothesis places no functional role per se, and rather argues that CMC results from the oscillatory nature of the sensorimotor activity (Bourguignon et al., 2017). In this view, a prominent ∼20-Hz rhythm sets the basis for modulations at ∼20-Hz in the motor command that is also seen in EMG activity and leads to coherence between cortical and muscle signals. Here, we will discuss how our results align (or not) with these hypotheses.

*Motor state maintenance* hypothesis. Reports of CMC being enhanced during isometric contraction, abolished during movement, and strong immediately post-movement (Baker et al., 1997; Kilner et al., 2003) have suggested a straightforward functional role for CMC: maintenance of the ongoing motor state. Increased CMC would act to preserve the motor action at the periphery, while decreased CMC would allow for motor flexibility. Distinct from previous reports, our data revealed the dynamic nature of CMC during an isometric contraction, on a time-scale of the order of 100 ms. A priori, the consideration of within-task modulation of CMC does not fit neatly into the above hypothesis. However, our data is perfectly in line with a dynamic version of the *motor state maintenance* hypothesis, but only if one embraces the view that steady contractions are highly dynamic processes (Taylor et al., 2003), where the motor state continuously commands either an increase or a decrease in force (those were indeed preceded by transiently increased CMC). In that setting, force plateaus indicate a change in the motor state that is subsequent to decreased CMC, as per the *motor state maintenance* hypothesis.

*Test-pulse* hypothesis. Here, CMC would capture the mechanism whereby ∼20-Hz ‘test pulses’ probe the periphery to allow for continuous motor recalibration (Mackay, 1997; Baker, 2007). A method that aims to assess the degree of synchronicity between oscillations in the SM1 cortex and muscles certainly appeals as a measurement of sensorimotor binding. Based on previous findings of phase-frequency dependence being incompatible with pure efferent origin of

CMC (Gerstner et al., 1996; Riddle and Baker, 2005; Baker, 2007) and on the existence of a significant effective coupling from EMG to SM1 activity (Tsujimoto et al., 2009; Witham et al., 2010, 2011; Lim et al., 2014), it has been proposed that CMC receives a contribution from both efferent and afferent signaling (for a review, see Bourguignon et al., 2019). If this hypothesis were true, CMC should modulate in response to changes in the periphery, including changes in the force, by virtue of the specialized sensory receptors (primarily the proprioceptors) located there. During sustained isometric contraction, this could be exemplified by increased CMC after fluctuations in force. However, our results revealed a temporal evolution of CMC not in line with this concept. Instead, we identified temporally consistent modulations in CMC across an analysis of the four force features, where changes in CMC always preceded the respective force component by ∼40 ms.

*SM1 rhythm corollary* hypothesis. Following this view, CMC arises because the corticospinal drive is modulated by the ∼20-Hz SM1 rhythm, and thus, is a byproduct of the oscillatory nature of population-level SM1 activity (Bourguignon et al., 2017). This is not to say that proprioceptive and sensory feedback cannot modify the sensorimotor rhythm, as afferent signaling must guide the motor command, only that such afferent signaling operates in a lower frequency channel (Bourguignon et al., 2017). This view is well supported by the tight temporal matching between CMC and power modulations. It is also compatible with the decoupling we observed between the ∼20-Hz and full-band EMG power. Indeed, because the ∼20-Hz sensorimotor rhythm shapes the corticospinal drive and the periphery receives this drive, it follows that ∼20-Hz EMG power emulates that of the cortex. Critically, decreases in ∼20-Hz SM1 power, indicative of sensorimotor involvement, can command reversal of force in either direction. This, in turn, leads to opposite changes in full-band EMG power, in relation to low and high force plateau signals (Fig. 5). The same reasoning applies to modulations in relation to force increase vs. decrease signals.

### Validation of methods through assessment of electromechanical coupling in muscles

The expected modulations in wide-band EMG power in relation to the force features validated our approach. About 12.5 ms before an increase or decrease in rate of force change, wide-band EMG power increased or decreased (respectively). This delay corresponds well with the expected electromechanical delay (time lag between onset of muscle activity and onset of force development), at least in active muscles where the muscle-tendon unit slack is readily taken up and only remains the contribution of the electrochemical component (excitation-contraction coupling of the muscle fibers) of the delay (Cavanagh and Komi, 1979). The wide-band EMG amplitude and muscle force output are highly correlated especially at low submaximal contraction intensities (De Luca, 1997). Therefore, it is unsurprising that a change in EMG power led to a change in contraction force in the same direction. The relationship between wide-band EMG power and force plateaus is not as well described in the literature. Force plateaus in the context of this study can be seen as the transition from an increase to decrease in rate of force (plateauing high), or vice versa (plateauing low). High plateaus were preceded (–2.0 ms) by a decrease in wide-band EMG power. This makes perfect sense: the force ceases to increase following a decrease in muscle activity. Conversely, low plateaus were preceded by an increase in power with a similar time course. Collectively, these results validate our novel weighting analysis method.

### Limitations and perspectives

Modulation in ∼20-Hz CMC, SM1 power and EMG power by specific force parameters occurred during a task that required very low force output from the hand muscles (2–4 N). Further studies would be needed to generalize these observations to higher intensity isometric contractions and dynamic contractions (concentric or eccentric). Indeed, nuanced changes in ∼20-Hz CMC and power do vary depending on the type and intensity of motor actions (Liu et al., 2019; Glories et al., 2021).

Future research will be necessary to expand our analysis approach while examining refined motor tasks requiring changes in rates of force, proprioceptive challenges, and additional contraction intensities. The continued study of neural force regulation is critical for insight into motor diseases as well as for the advancement of assistive technologies. Along these lines, whether CMC and sensorimotor power modulation may be exploited for intervention remains to be elucidated.

## Conclusion

Our study provides insight into the dynamics of sensorimotor oscillations at play in the maintenance of steady muscle contractions. Using a novel analysis approach, which we validated through assessment of muscle electromechanical coupling, we have demonstrated hitherto unfathomable short-lived modulations of ∼20-Hz CMC and SM1 oscillations in relation to force fluctuations. Our results have implications for the existing theories regarding the functional role of CMC and for the role of the SM1 cortex in sustaining muscle contractions: steady contractions entail stable rather than fluttering SM1 cortex involvement.

## Contributions

H.P., V.J., M.B. designed the study; H.P., M.B. collected the data; S.J.M., T.L., M.V.G, G.N., M.B. analyzed the data; S.J.M. wrote the initial version of the manuscript; and all authors discussed the results and their interpretation, and reviewed and edited the manuscript.

## Acknowledgments

Scott Mongold, Thomas Legrand, and Mathieu Bourguignon were supported by the Fonds de la Recherche Scientifique (F.R.S.-FNRS, Brussels, Belgium; grant MIS F.4504.21). Harri Piitulainen was supported by the Academy of Finland (grants 266133, 296240, 326988, 327288 and 311877) including ”Brain changes across the life-span” profiling funding to University of Jyväskylä.

We thank Helge Kainulainen and Ronny Schreiber at Aalto NeuroImaging for providing technical help and the force sensor system for the study.

We thank Riitta Hari for her participation in the initial study.

## References

1. Androulidakis AG, Doyle LMF, Yarrow K, Litvak V, Gilbertson TP, Brown P (2007) Anticipatory changes in beta synchrony in the human corticospinal system and associated improvements in task performance. Eur J Neurosci 25:3758–3765.

2. Aumann TD, Prut Y (2015) Do sensorimotor β-oscillations maintain muscle synergy representations in primary motor cortex? Trends Neurosci 38:77–85.

3. Averbeck BB, Lee D (2004) Coding and transmission of information by neural ensembles. Trends Neurosci 27:225–230.

4. Baker SN (2007) Oscillatory interactions between sensorimotor cortex and the periphery. Curr Opin Neurobiol 17:649–655.

5. Baker SN, Olivier E, Lemon RN (1997) Coherent oscillations in monkey motor cortex and hand muscle EMG show task-dependent modulation. J Physiol 501 ( Pt 1):225–241.

6. Bayraktaroglu Z, von Carlowitz-Ghori K, Curio G, Nikulin VV (2013) It is not all about phase: Amplitude dynamics in corticomuscular interactions. NeuroImage 64:496–504 Available at: http://dx.doi.org/10.1016/j.neuroimage.2012.08.069.

7. Bishara AJ, Hittner JB (2015) Reducing Bias and Error in the Correlation Coefficient Due to Nonnormality. Educ Psychol Meas 75:785–804.

8. Bourguignon M, Jousmäki V, Dalal SS, Jerbi K, De Tiège X (2019) Coupling between human brain activity and body movements: Insights from non-invasive electromagnetic recordings. Neuroimage 203:116177.

9. Bourguignon M, Piitulainen H, De Tiège X, Jousmäki V, Hari R (2015) Corticokinematic coherence mainly reflects movement-induced proprioceptive feedback. NeuroImage 106:382–390 Available at: http://dx.doi.org/10.1016/j.neuroimage.2014.11.026.

10. Bourguignon M, Piitulainen H, Smeds E, Zhou G, Jousmäki V, Hari R (2017) MEG Insight into the Spectral Dynamics Underlying Steady Isometric Muscle Contraction. J Neurosci 37:10421–10437.

11. Cavanagh PR, Komi PV (1979) Electromechanical delay in human skeletal muscle under concentric and eccentric contractions. Eur J Appl Physiol Occup Physiol 42:159–163.

12. Conway BA, Halliday DM, Farmer SF, Shahani U, Maas P, Weir AI, Rosenberg JR (1995) Synchronization between motor cortex and spinal motoneuronal pool during the performance of a maintained motor task in man. J Physiol 489 ( Pt 3):917–924.

13. Danion F, Galléa C (2004) The relation between force magnitude, force steadiness, and muscle co-contraction in the thumb during precision grip. Neurosci Lett 368:176–180.

14. De Luca CJ (1997) The Use of Surface Electromyography in Biomechanics. Journal of Applied Biomechanics 13:135–163 Available at: http://dx.doi.org/10.1123/jab.13.2.135.

15. Démas J, Bourguignon M, Périvier M, De Tiège X, Dinomais M, Van Bogaert P (2019) Mu rhythm: State of the art with special focus on cerebral palsy. Ann Phys Rehabil Med Available at: http://dx.doi.org/10.1016/j.rehab.2019.06.007.

16. Efron B (1979) Bootstrap Methods: Another Look at the Jackknife. The Annals of Statistics 7 Available at: http://dx.doi.org/10.1214/aos/1176344552.

17. Efron B, Tibshirani RJ (1994) An Introduction to the Bootstrap. Chapman and Hall.

18. Engel AK, Fries P (2010) Beta-band oscillations—signalling the status quo? Current Opinion in Neurobiology 20:156–165 Available at: http://dx.doi.org/10.1016/j.conb.2010.02.015.

19. Enoka RM, Farina D (2021) Force Steadiness: From Motor Units to Voluntary Actions. Physiology 36:114–130.

20. Faes L, Pinna GD, Porta A, Maestri R, Nollo GD (2004) Surrogate Data Analysis for Assessing the Significance of the Coherence Function. IEEE Transactions on Biomedical Engineering 51:1156–1166 Available at: http://dx.doi.org/10.1109/tbme.2004.827271.

21. Fisher RJ, Galea MP, Brown P, Lemon RN (2002) Digital nerve anaesthesia decreases EMG-EMG coherence in a human precision grip task. Experimental Brain Research 145:207–214 Available at: http://dx.doi.org/10.1007/s00221-002-1113-x.

22. Gerstner W, van Hemmen JL, Cowan JD (1996) What matters in neuronal locking? Neural Comput 8:1653–1676.

23. Gilbertson T (2005) Existing Motor State Is Favored at the Expense of New Movement during 13-35 Hz Oscillatory Synchrony in the Human Corticospinal System. Journal of Neuroscience 25:7771–7779 Available at: http://dx.doi.org/10.1523/jneurosci.1762-05.2005.

24. Glories D, Soulhol M, Amarantini D, Duclay J (2021) Specific modulation of corticomuscular coherence during submaximal voluntary isometric, shortening and lengthening contractions. Sci Rep 11:6322.

25. Gross J, Timmermann L, Kujala J, Dirks M, Schmitz F, Salmelin R, Schnitzler A (2002) The neural basis of intermittent motor control in humans. Proc Natl Acad Sci U S A 99:2299–2302.

26. Halliday DM, Rosenberg JR, Amjad AM, Breeze P, Conway BA, Farmer SF (1995) A framework for the analysis of mixed time series/point process data—Theory and application to the study of physiological tremor, single motor unit discharges and electromyograms. Progress in Biophysics and Molecular Biology 64:237–278 Available at: http://dx.doi.org/10.1016/s0079-6107(96)00009-0.

27. Hari R, Bourguignon M, Piitulainen H, Smeds E, De Tiège X, Jousmäki V (2014) Human primary motor cortex is both activated and stabilized during observation of other person’s phasic motor actions. Philos Trans R Soc Lond B Biol Sci 369:20130171.

28. Hari R, Salenius S (1999) Rhythmical corticomotor communication. Neuroreport 10:R1–R10.

29. Jones KE, Hamilton AF, Wolpert DM (2002) Sources of signal-dependent noise during isometric force production. J Neurophysiol 88:1533–1544.

30. Kilner JM, Baker SN, Salenius S, Hari R, Lemon RN (2000) Human cortical muscle coherence is directly related to specific motor parameters. J Neurosci 20:8838–8845.

31. Kilner JM, Baker SN, Salenius S, Jousmäki V, Hari R, Lemon RN (1999) Task-dependent modulation of 15-30 Hz coherence between rectified EMGs from human hand and forearm muscles. J Physiol 516 ( Pt 2):559–570.

32. Kilner JM, Fisher RJ, Lemon RN (2004) Coupling of Oscillatory Activity Between Muscles Is Strikingly Reduced in a Deafferented Subject Compared With Normal Controls. Journal of Neurophysiology 92:790–796 Available at: http://dx.doi.org/10.1152/jn.01247.2003.

33. Kilner JM, Salenius S, Baker SN, Jackson A, Hari R, Lemon RN (2003) Task-dependent modulations of cortical oscillatory activity in human subjects during a bimanual precision grip task. Neuroimage 18:67–73.

34. Kristeva R, Patino L, Omlor W (2007) Beta-range cortical motor spectral power and corticomuscular coherence as a mechanism for effective corticospinal interaction during steady-state motor output. Neuroimage 36:785–792.

35. Laidlaw DH, Bilodeau M, Enoka RM (2000) Steadiness is reduced and motor unit discharge is more variable in old adults. Muscle Nerve 23:600–612.

36. Lim M, Kim JS, Kim M, Chung CK (2014) Ascending beta oscillation from finger muscle to sensorimotor cortex contributes to enhanced steady-state isometric contraction in humans. Clin Neurophysiol 125:2036–2045.

37. Liu J, Sheng Y, Liu H (2019) Corticomuscular Coherence and Its Applications: A Review. Front Hum Neurosci 13:100.

38. Mackay WA (1997) Synchronized neuronal oscillations and their role in motor processes. Trends Cogn Sci 1:176–183.

39. McAuley JH, Rothwell JC, Marsden CD (1997) Frequency peaks of tremor, muscle vibration and electromyographic activity at 10 Hz, 20 Hz and 40 Hz during human finger muscle contraction may reflect rhythmicities of central neural firing. Exp Brain Res 114:525–541.

40. Mendez-Balbuena I, Huethe F, Schulte-Mönting J, Leonhart R, Manjarrez E, Kristeva R (2012) Corticomuscular coherence reflects interindividual differences in the state of the corticomuscular network during low-level static and dynamic forces. Cereb Cortex 22:628–638.

41. Missenard O, Mottet D, Perrey S (2009) Factors responsible for force steadiness impairment with fatigue. Muscle Nerve 40:1019–1032.

42. Muthukumaraswamy SD, Singh KD (2011) A cautionary note on the interpretation of phase-locking estimates with concurrent changes in power. Clin Neurophysiol 122:2324–2325.

43. Nichols TE, Holmes AP (2002) Nonparametric permutation tests for functional neuroimaging: a primer with examples. Hum Brain Mapp 15:1–25.

44. Oldfield RC (1971) The assessment and analysis of handedness: The Edinburgh inventory. Neuropsychologia 9:97–113 Available at: http://dx.doi.org/10.1016/0028-3932(71)90067-4.

45. Piitulainen H, Botter A, Bourguignon M, Jousmäki V, Hari R (2015a) Spatial variability in cortex-muscle coherence investigated with magnetoencephalography and high-density surface electromyography. J Neurophysiol 114:2843–2853.

46. Piitulainen H, Bourguignon M, Smeds E, De Tiège X, Jousmäki V, Hari R (2015b) Phasic stabilization of motor output after auditory and visual distractors. Hum Brain Mapp 36:5168–5182.

47. Pineda JA (2005) The functional significance of mu rhythms: translating “seeing” and “hearing” into “doing.” Brain Res Brain Res Rev 50:57–68.

48. Purves D (1999) Neurosciences. De Boeck.

49. Riddle CN, Baker SN (2005) Manipulation of peripheral neural feedback loops alters human corticomuscular coherence. J Physiol 566:625–639.

50. Salenius S, Hari R (2003) Synchronous cortical oscillatory activity during motor action. Curr Opin Neurobiol 13:678–684.

51. Salenius S, Portin K, Kajola M, Salmelin R, Hari R (1997) Cortical control of human motoneuron firing during isometric contraction. J Neurophysiol 77:3401–3405.

52. Steiger JH (1980) Tests for comparing elements of a correlation matrix. Psychol Bull 87:245–251.

53. Taylor AM, Christou EA, Enoka RM (2003) Multiple features of motor-unit activity influence force fluctuations during isometric contractions. J Neurophysiol 90:1350–1361.

54. Theiler J, Eubank S, Longtin A, Galdrikian B, Doyne Farmer J (1992) Testing for nonlinearity in time series: the method of surrogate data. Physica D: Nonlinear Phenomena 58:77–94 Available at: http://dx.doi.org/10.1016/0167-2789(92)90102-s.

55. Thomson DJ (1982) Spectrum estimation and harmonic analysis. Proceedings of the IEEE 70:1055–1096 Available at: http://dx.doi.org/10.1109/proc.1982.12433.

56. Tracy BL, Enoka RM (2002) Older adults are less steady during submaximal isometric contractions with the knee extensor muscles. J Appl Physiol 92:1004–1012.

57. Tsujimoto T, Mima T, Shimazu H, Isomura Y (2009) Directional organization of sensorimotor oscillatory activity related to the electromyogram in the monkey. Clin Neurophysiol 120:1168–1173.

58. Ushiyama J, Katsu M, Masakado Y, Kimura A, Liu M, Ushiba J (2011a) Muscle fatigue-induced enhancement of corticomuscular coherence following sustained submaximal isometric contraction of the tibialis anterior muscle. J Appl Physiol 110:1233–1240.

59. Ushiyama J, Suzuki T, Masakado Y, Hase K, Kimura A, Liu M, Ushiba J (2011b) Between-subject variance in the magnitude of corticomuscular coherence during tonic isometric contraction of the tibialis anterior muscle in healthy young adults. J Neurophysiol 106:1379–1388.

60. Ushiyama J, Takahashi Y, Ushiba J (2010) Muscle dependency of corticomuscular coherence in upper and lower limb muscles and training-related alterations in ballet dancers and weightlifters. J Appl Physiol 109:1086–1095.

61. Ushiyama J, Yamada J, Liu M, Ushiba J (2017) Individual difference in β-band corticomuscular coherence and its relation to force steadiness during isometric voluntary ankle dorsiflexion in healthy humans. Clin Neurophysiol 128:303–311.

62. Witham CL, Riddle CN, Baker MR, Baker SN (2011) Contributions of descending and ascending pathways to corticomuscular coherence in humans. J Physiol 589:3789–3800.

63. Witham CL, Wang M, Baker SN (2010) Corticomuscular coherence between motor cortex, somatosensory areas and forearm muscles in the monkey. Front Syst Neurosci 4 Available at: http://dx.doi.org/10.3389/fnsys.2010.00038.

64. Witte M, Patino L, Andrykiewicz A, Hepp-Reymond M-C, Kristeva R (2007) Modulation of human corticomuscular beta-range coherence with low-level static forces. Eur J Neurosci 26:3564–3570.

